# Slow-oscillatory tACS does not modulate human motor cortical response to repeated plasticity paradigms

**DOI:** 10.1101/2021.09.20.461168

**Authors:** Claire Bradley, Jessica Elliott, Samuel Dudley, Genevieve A. Kieseker, Jason B. Mattingley, Martin V. Sale

## Abstract

Previous history of activity and learning modulates synaptic plasticity and can lead to saturation of synaptic connections. According to the synaptic homeostasis hypothesis, neural oscillations during slow-wave sleep play an important role in restoring plasticity within a functional range. However, it is not known whether slow-wave oscillations – without the concomitant requirement of sleep – play a causal role in human synaptic homeostasis. Here, slow-oscillatory transcranial alternating current stimulation (tACS, 1Hz, 1mA, 18 minutes) was interleaved between two plasticity-inducing interventions: motor learning, and a paradigm known to induce long-term-potentiation-like plasticity in human motor cortex (paired associative stimulation; PAS). The hypothesis tested was that slow-oscillatory tACS would abolish the expected interference between motor learning and PAS, and facilitate plasticity from successive interventions. Thirty-six participants received sham and active fronto-motor tACS in two separate sessions, along with electroencephalography (EEG) recordings. A further 38 participants received tACS through a control (posterior midline) montage. Using neuro-navigated transcranial magnetic stimulation (TMS) over the left motor cortex, motor evoked potentials (MEPs) were recorded throughout the session. Bayesian statistics were used to quantify evidence for or against the hypothesis of an effect of each intervention on MEP amplitude. As expected, there was converging evidence that motor training increased MEPs. Importantly, we found moderate evidence *against* an effect of active tACS in restoring PAS plasticity, and no evidence of lasting entrainment of slow-oscillations in the EEG. This suggests that, under the conditions tested here, slow-oscillatory tACS does not modulate synaptic homeostasis in the motor system of awake humans.

## Introduction

Synaptic plasticity – the modulation of synaptic neural transmission strength – is a prominent biological mechanism underlying learning (Moser et al., 1998; Rioult-Pedotti et al., 2000; Hodgson et al., 2005). Importantly, the likelihood that a synapse will undergo plastic change is not constant over time, but depends on previous history of synaptic activity and plasticity – a phenomenon known as ‘meta-plasticity’ (Abraham and Bear, 1996). Reflective of this, interventions known to trigger plasticity in the human motor system interfere with, and even reverse, each other’s effects when applied sequentially (Shadmehr and Brashers-Krug, 1997; Ziemann et al., 2004, 2008), highlighting the importance of previous history of network activity for plasticity effects.

Sleep is a major physiological modulator of learning and plasticity (Maquet, 2001; Diekelmann and Born, 2010). Whether for a whole night or a short nap, sleep has been shown to enhance retention of learned material (Tucker et al., 2006; Mascetti et al., 2013) and allows further acquisition (Mander et al., 2011; Antonenko et al., 2013). At a synaptic level, sleep has been hypothesized to achieve a general downscaling of synaptic weights (Tononi and Cirelli, 2014). This account of sleep function – the synaptic homeostasis hypothesis – posits that synapse modifications accumulate during the waking day due to learning and non-specific exposure to environmental stimuli. The resulting increased synaptic strength imposes metabolic and computational pressure, which is resolved through re-normalization of synaptic strength during sleep. Although experimental studies testing this hypothesis in humans are scarce, it appears sleep may recalibrate cortical excitability (Huber et al., 2013), synaptic plasticity (Kuhn et al., 2016), and learning (Fattinger et al., 2017) in healthy adults.

Slow-wave oscillations – low frequency oscillatory electrical activity characteristic of non-rapid eye movement sleep – is a prime candidate mechanism for synaptic strength downscaling. Slow-wave oscillations increase after a learning episode (Mölle et al., 2004) and with time awake, whereas they decrease during sleep as the night progresses (Tononi and Cirelli, 2014). Boosting their amplitude and duration during sleep by means of non-invasive brain stimulation results in better encoding of information and increased memory retention (Marshall et al., 2006; Antonenko et al., 2013; Ladenbauer et al., 2016, 2017), whereas disrupting them impairs the restorative property of sleep (Aeschbach et al., 2008; Fattinger et al., 2017; although see Paßmann et al., 2016). Interestingly, a causal link between slow-wave oscillations and learning is further suggested by the finding that these oscillations may be entrained in the *awake* human brain and that such entrainment can enhance encoding processes (Kirov et al., 2009). However, whether slow-wave oscillations – without the concomitant requirement of sleep – play a causal role in synaptic homeostasis has never been investigated in humans.

Here, we investigated whether slow-oscillatory activity, in line with its hypothesized synaptic downscaling function, enables successive plasticity interventions to exert their full effects. More specifically, we asked whether a short episode of non-invasive brain stimulation known as slow-oscillatory transcranial alternating current stimulation (tACS), comparable in length to a nap, can attenuate the interference between two successive motor plasticity paradigms in awake humans. In 36 participants, we interposed 18 minutes’ of tACS (either active or sham) between two interventions known to induce plastic changes in the motor cortex: (1) 30 minutes of ballistic thumb movement (motor training, MT), and (2) excitatory paired associative stimulation (PAS) (**Fig. 1**). We quantified motor cortical plasticity indirectly by measuring the amplitude of transcranial magnetic stimulation (TMS)-evoked motor evoked potentials (MEPs). Consistent with the literature, we hypothesized that motor training would lead to an increase in the amplitude of MEPs (Muellbacher et al., 2001, 2002; Ziemann et al., 2004), that this increase would persist after sham tACS (Muellbacher et al., 2001; Ziemann et al., 2004), and that it would cause the subsequent PAS paradigm to be ineffective (Ziemann et al., 2004). In contrast, we hypothesized that following active tACS, and in spite of the previous motor training, PAS would result in an increase in MEPs, consistent with a synaptic downscaling effect as predicted by the synaptic homeostasis hypothesis. In a further 38 participants, we delivered active tACS to a control montage that does not target motor networks, hypothesizing that the effect of control tACS would be similar to sham. We used Bayesian statistics to quantify the extent of evidence for or against our various hypotheses. This approach is preferable to null-hypothesis significance testing, in that it can distinguish between situations where there is evidence for the null hypothesis, for the alternative hypothesis or not enough evidence to distinguish between both. This can be particularly useful in interpreting the absence of difference between two conditions but applies more generally to all inferences (Biel & Friedrich 2018; Dienes & Maclatchie 2018).

**Figure 1.**
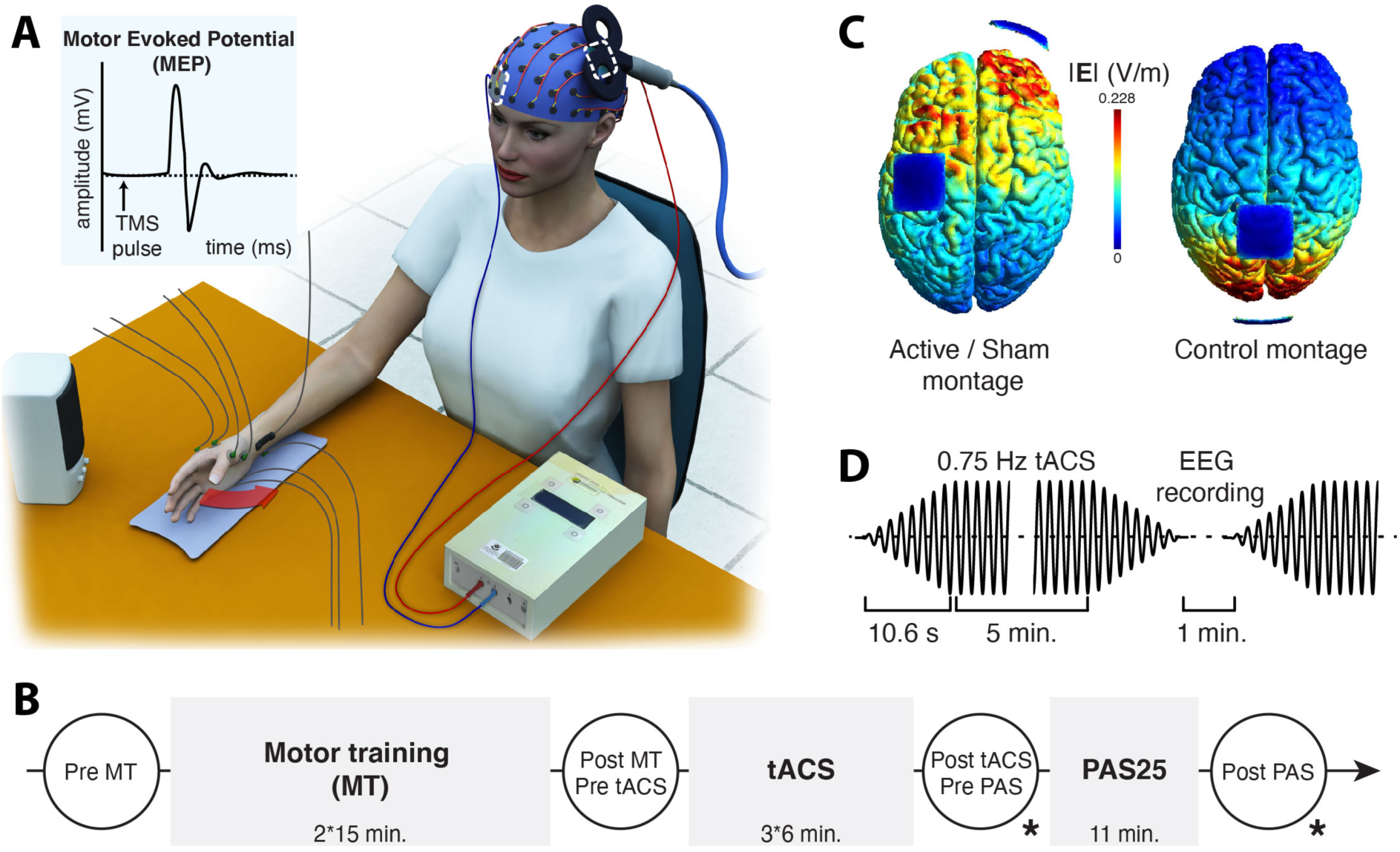
Experimental setup and study protocol. **A)** In the active vs. sham condition, transcranial magnetic stimulation (TMS) was combined with electromyography to record motor evoked potentials (MEPs, i.e. the muscle response to a TMS pulse, *blue inset*) from right *abductor pollicis brevis* (APB) and two other intrinsic hand muscles (first dorsal interosseous, FDI and *abductor digiti minimi*, ADM). The TMS coil was positioned over left primary motor cortex, on top of an electroencephalography (EEG) cap (used to record electrical brain activity) and a transcranial alternating current stimulation (tACS) electrode (used to deliver electrical stimulation, *dashed white lines*). The return electrode for tACS was placed over the contralateral supra-orbit. A peripheral nerve stimulator was placed on the right wrist of participants to allow median nerve stimulation for the paired associative stimulation protocol. An accelerometer was taped to the participant’s right thumb to measure thumb speed during motor training (*red arrow*, accelerometer not shown here). Setup was identical in the control tACS condition, except that a different tACS montage was used and there was no recording of FDI, ADM or EEG. **B)** Session timeline of MEP recordings (white circles) and three successive experimental manipulations (grey rectangles). MT: motor training, tACS: transcranial alternating current stimulation, PAS: paired associative stimulation. **\*** denotes additional MEPs collected at an adjusted stimulation intensity, in a subset of participants. **C)** Cortical grey matter projection of the norm of the electric field (V/m) generated by the active vs. sham montage (left) and the control montage (right). Dark blue squares represent the two tACS electrodes. Modeling and visualization were performed in SimNIBS (Thielscher et al. 2015). **D)** Summary stimulus waveform for one out of three segments of tACS stimulation. In the active vs. sham condition, resting EEG was recorded continuously from 1 minute before the start of tACS to one minute after the end of tACS, allowing extraction of 1-minute periods prior to and immediately following each tACS segment. Sham stimulation only consisted of the ramp-up and ramp-down, after which stimulation was turned off (total stimulation time: 63.6s).

## Materials and Methods

### Participants

Seventy-eight participants (mean age ± SD: 24 ± 5 years; 50 women; split into two groups) took part in the study. All participants were right-handed (mean laterality quotient = 0.92, range 0.5-1.00) as assessed by the Edinburgh Handedness Questionnaire (Oldfield, 1971). All participants gave written informed consent prior to participation in the study, which was approved by The University of Queensland Human Research Ethics Committee. Participants were screened for family history of epilepsy, consumption of neuroactive drugs, and history of neurosurgery or brain injury, using a TMS safety questionnaire (Keel et al., 2000). Four participants were excluded from the final sample due to non-compliance with motor training (n=1), a request to exit the study before the end of the last session (n=1), very small MEPs (<0.1mV, n=1), and tDCS equipment failure (n=1). Final participant numbers were as follows. Active vs. sham tACS condition: n=36, mean age ± SD: 24 ± 4 years; 22 women; mean laterality quotient = 0.94. Control tACS montage condition: n=38, mean age ± SD: 24 ± 5 years; 28 women; mean laterality quotient = 0.89.

### Overview of experimental procedure

In the active vs. sham tACS condition, participants attended two experimental sessions, separated by at least one week. After general experimental setup, the cortical motor hotspot was located on the scalp, and motor and peripheral electrical thresholds were measured, as described in detail below. Participants then performed a motor training (MT) task for 30 minutes (**Fig. 1A, 1B**), followed by 18 minutes of tACS (fronto-motor montage), after which a further period of plasticity was induced in the motor cortex using TMS and concurrent stimulation of the median nerve (PAS protocol). MEPs and resting EEG recordings were collected before motor training, before tACS, as well as before and after PAS plasticity induction. Resting motor threshold was re-evaluated after PAS to investigate possible cellular/intrinsic plasticity effects (Delvendahl et al., 2012). The two sessions were identical except that in one of them, sham (placebo) tACS was delivered instead of active stimulation. Delivery of sham and active tACS was randomized and counter-balanced across participants, and double-blinded. For each participant, the two experiments were performed at the same time of day to minimise response variability due to circadian factors (Sale et al., 2007, 2008). A similar procedure was followed in the control tACS montage condition, as described in detail below.

### Neuro-navigated transcranial magnetic stimulation (TMS) of left motor cortex

Monophasic TMS pulses were applied through a figure-of-eight coil (outer diameter of each wing 70 mm) connected to a Magstim 200^2^ magnetic stimulator (Magstim, UK). The coil was held tangentially to the skull with the handle pointing backwards and laterally at an angle of 45° to the sagittal plane, at the optimal scalp site to evoke an MEP in the relaxed *abductor pollicis brevis* (APB) muscle of the right hand. With this coil placement, current flow was induced in a posterior-to-anterior direction in the brain. A neuro-navigation system (Visor2, ANT Neuro, The Netherlands) that tracks the position of the coil relative to the participant’s head ensured consistent placement of the TMS coil throughout the entire session (ensuring deviations of no more than 5 mm and 5° error in position and angle).

### Assessing plasticity in the cortico-spinal tract: motor-evoked potentials (MEPs)

Participants were seated comfortably in an experimental chair with their arms resting on a table and their eyes open, gazing at a fixation cross on a screen in front of them (**Fig. 1A**). Surface electromyographic (EMG) recordings from the APB, first dorsal interosseous (FDI) and *abductor digiti minimi* (ADM) muscles of the right hand were obtained using bipolar Ag-AgCl electrodes in a belly-tendon montage. EMG signals were amplified 1000 times, filtered (20 Hz – 2000 Hz, 50Hz notch filter) via a NeuroLog system (Digitimer, UK), digitized online with a data acquisition interface and Signal software (CED, Cambridge Electronic Design, UK) and stored on computer for offline analysis. The EMG signal from the APB muscle was continuously monitored on an oscilloscope throughout the session to ensure that the muscle was relaxed (absence of background activity, which would otherwise bias MEP measurements).

Resting motor threshold was defined as the intensity eliciting MEPs above 50 μV in 5 out of 10 consecutive trials (Rossini et al., 1994). Resting motor threshold was assessed at baseline and once again just after PAS. Mean threshold ± SD at baseline and after PAS was 52% ± 8% and 52.5% ± 8% of maximum stimulator output in the active session, and 52% ± 9% and 52% ± 10% in the sham tACS session. Test intensity was set at 130% of resting motor threshold. Cortico-spinal excitability was assessed at several time points during each experimental session: before motor training, 5 minutes after motor training (e.g. immediately before tACS), immediately after tACS and 5 minutes after PAS. Mean peak-to-peak amplitude of the APB MEP at rest was calculated by averaging the individual peak-to-peak amplitudes of MEPs elicited by 21 separate TMS pulses (delivered at ∼0.2 Hz; first pulse systematically discarded to avoid startle responses). In a subset of participants (n=29), TMS stimulation intensity post-tACS (pre-PAS) was adjusted so that MEPs had an amplitude that matched baseline MEPs (pre-motor training). If this amplitude adjustment took more than a few minutes, the second MEP block was recorded at the standard intensity (130% of resting motor threshold), so the timing of MEP assessments was not jeopardised. This intensity-adjusted MEP measure was taken again after PAS, in a counter-balanced order with the original intensity.

### Motor training

The motor training task was adapted from Muellbacher et al. (2001) and Ziemann et al. (2004). Participants’ right arm rested on a table, bent 90° at the elbow and attached to a guide to ensure consistent initial position for the movement. Participants were requested to perform their fastest possible thumb abduction movements of the right hand for 30 min (2*15 minutes with one minute break), paced by a tone at a rate of 0.5 Hz. After a brief familiarisation with the task and movement, acceleration was measured using a custom-made accelerometer mounted on the thumb; trial-by-trial acceleration in the horizontal plane was displayed on a monitor and participants were repeatedly encouraged to maximize this component. The peak-to-peak amplitude of acceleration in the horizontal plane for the first 20 and the last 20 trials (excluding familiarisation period) were averaged to quantify performance change following motor training.

### Slow-oscillatory transcranial alternating current stimulation (tACS)

tACS was administered via a NeuroConn stimulator (neuroCare Group GmbH, Germany) with two 4 × 4 cm electrodes. One electrode was positioned over the motor cortical hotspot, the other electrode was placed over the right supraorbital region (**Fig. 1C**). Electrodes were held in place by conductive paste (Ten20 EEG paste, Weaver and company, USA). tACS was applied at a stimulus intensity of 1 mA (resulting in a current density of 0.0625 mA/cm^2^) and a frequency of 0.75 Hz. Stimulation was applied over 3 x five-minute blocks, each separated by 1 minute (to allow for EEG recording, and in keeping with Marshall et al. (2006) and Kirov et al. (2009)). Each stimulation block consisted of 225 cycles (300 seconds) and included a ramp up and ramp down period of 8 cycles (∼10 s) (**Fig. 1D**). For the sham condition, the stimulus parameters were the same except that following the 8-cycle ramp-up and ramp-down, the stimulator was turned off. Participants were not informed about the type of stimulation they were receiving; they sat in silence with their eyes open.

### Paired associative stimulation (PAS)

The PAS paradigm involved a series of paired peripheral and cortical stimuli (Stefan et al., 2000; Wolters et al., 2003; Ziemann et al., 2004). A peripheral electrical stimulus was delivered to the median nerve of the right wrist at an intensity that evoked a clear response in the APB muscle of at least 0.2 mV amplitude (Active vs. sham tACS condition: mean ± SD perceptual threshold: 3.17 ± 0.8 mA, stimulation intensity: 7.1 ± 2.3 mA; Control tACS montage condition: mean ± SD perceptual threshold: 2 ± 0.8 mA, stimulation intensity: 7.4 ± 2.5 mA), using a constant current stimulator (DS7 stimulator; Digitimer, UK) with bipolar surface electrodes separated by 30 mm (cathode proximal). Stimuli were square waves with a pulse width of 200 μs. Each electrical stimulus was followed by suprathreshold TMS (intensity: 130% of resting motor threshold) over the hand area of the contralateral (left) motor cortex, with a 25 ms delay. A total of 132 paired peripheral and cortical stimuli were delivered at an average frequency of 0.2 Hz. Because spatial attention has been shown to enhance the effects of PAS (Kamke et al., 2014), participants were asked to overtly monitor a blinking LED strapped to their right thumb for infrequent changes in the rhythm of blinking and to report the number of events at the end of the PAS stimulation.

### Electroencephalography (EEG) recordings of brain activity

To investigate potential entrainment of endogenous slow oscillations by active tACS (Kirov et al., 2009), 64-channel EEG was recorded at rest with eyes open in short 1-minute continuous segments before and just after each block of tACS (active vs. sham conditions only). EEG could not be collected in three participants due to the EEG cap not fitting (n=1), not enough time to apply the cap in one session (n=1) and motor threshold being too high with the cap on (n=1).

#### EEG recording

Spontaneous EEG was recorded with a TMS-compatible 64-channel EEG cap (BrainCap, BrainProducts, Germany; Ag/AgCl electrodes) in accordance with the 10–10 extended international system. All electrodes were referred to the right mastoid and impedance was kept below 5 kΩ using a viscous electrode paste (Abralyt HiCl Gel, EasyCap, Germany); the ground electrode was incorporated in the cap at AFz. BrainRecorder software and BrainAmp MR Plus amplifiers (BrainProducts, Germany) with a 5000 Hz sampling rate and low-pass filter of 1000 Hz (no high-pass filter, resulting in a DC recording) were used to record periods of EEG throughout the experiment. Participants were at rest, with eyes open and gaze steadied on a fixation point.

#### EEG analysis

Four segments of 1-minute resting EEG were extracted from the continuous recordings: one before tACS and one after each 5-minute block of tACS stimulation, using BrainVision Analyser (Brain Products, Germany). Special care was taken that segments started no later than 10ms after the end of the active stimulation (as evidenced by a clear artefact); segments in the sham condition were taken at the same time-points as in the active condition. Some electrodes presented an exponential decaying artefact following active tACS termination in most participants; these electrodes were discarded from further analysis (9 electrodes in total, all located near or on the tACS stimulation pads: FC3, FC1, C3, C1, CP1, CP3, Fp2, AF4, AF8), as were TP9 and TP10 (systematically noisy) and Fp1 and Fpz (because of TMS neuronavigation markers). Data were imported in EEGLAB v13.6.5b (Delorme & Makeig, 2004) running in Matlab R2016a, segmented, down-sampled to 1000Hz and filtered (notch: 50Hz, low-pass: 100Hz). Periods of noise (visual inspection for speech, movements) were discarded; similarly, noisy or flat electrodes were interpolated. ICA analysis (extended binICA algorithm) was performed on filtered (band-pass:2-35Hz), PCA-treated data (to compensate for rank-reduction induced by interpolating electrodes), in order to identify and reject vertical and horizontal eye movement artefacts (Chaumon, Bishop, & Busch, 2015). The correction (ICA weights and sphering matrix) was then applied to the less-filtered data that were considered for further analysis. The power spectrum density was calculated using the ‘spectopo’ function of EEGLAB (average of FFT of 5s segments, Hanning-window, 50% overlap). Average power values were taken over 6 contiguous frequency bands (as in Kirov, Weiss, Siebner, Born, & Marshall, 2009): tACS-frequency: [0.5-1Hz], delta: [1-4Hz], theta: [4-8Hz], low alpha: [8-12Hz], high alpha: [12-15Hz] and beta: [15-25Hz]. Values were further averaged over the three time-points post-tACS and decibel-change values relative to baseline (pre-tACS) were computed.

#### EEG statistical analysis

Of the thirty-three participants with EEG recordings, eight participants were excluded due to poor data quality as assessed during the artefact rejection phase, resulting in a final sample of n=25 participants. Cluster-based permutation testing was performed in FieldTrip (Oostenveld, Fries, Maris, & Schoffelen, 2011) (sample statistic: dependent samples t-test; test statistic: maximum of sum of t-values, MonteCarlo method, 1000 permutations, cluster-forming alpha level: 0.05, two-tailed test alpha level: 0.05) in order to compare: (a) baseline maps before active and sham tACS, (b) post-tACS to pre-tACS (active and sham separately), (c) decibel (dB)-change post-active tACS to post-sham tACS. Signal from the 10 electrodes surrounding the motor tACS pad (FCz, Cz, CPz, CP5, C5, FC5, F3, F1, P1, P3) was pooled to extract mean power values in the tACS frequency over all time-points (Pre, Post 5’, Post 10’ and Post 15’ tACS). These were analysed in GraphPad Prism7, by means of a two-way repeated measures ANOVA, with factors Time (Pre, Post 5’, Post 10’ and Post 15’) and tACS type (Active and Sham), followed by a post-hoc Dunnett’s multiple comparisons test, comparing each ‘Post’ time-point to baseline (‘Pre’).

### Control tACS condition: posterior midline montage

Participants in the control tACS condition underwent procedures identical to those described above, except for the following differences. Participants attended only one experimental session, with no EEG recording. MEPs were recorded from the APB muscle only. Resting motor threshold was defined as the intensity eliciting MEPs above 50 μV in 10 out of 20 consecutive trials (Rossini et al. 2015), and MEPs were calculated from the average of 41 separate pulses instead of 21. Finally, intensity was not adjusted to 1 mV before PAS and motor threshold was not re-measured after PAS, resulting in a comparable number of pulses delivered throughout the session in all groups. Mean resting motor threshold ± SD at baseline was 46% ± 10% of maximum stimulator output. Importantly, control (active) tACS was delivered through a posterior midline montage (Oz-CPz, see **Fig. 1D** for an electric field distribution). This montage was chosen to minimize overlap between its electric field and that of the fronto-motor active montage, and with areas supporting motor learning in general (including the cerebellum).

### Statistical analysis

Statistical analyses were conducted in JASP (JASP team, 2020). Data analysed were acceleration and raw MEP amplitude (as recommended by Lahr et al. 2016). In the active vs sham condition (n=36), Bayesian two-way repeated-measures ANOVAs, with factors ‘time’ (‘Pre’ and ‘Post’ intervention) and ‘tACS type’ (‘active’ or ‘sham’ tACS) were used to analyse acceleration and MEP amplitude data. Four ANOVAs were conducted: one to examine the effect of motor training and tACS type on acceleration data, and three separate ANOVAs to examine the effect of the three interventions (motor training, tACS and PAS) and tACS type on MEP amplitude. Post-hoc Bayesian paired t-tests were used to explore specific contrasts of interest. In the control tACS montage condition (N=38), Bayesian paired t-tests were used to examine the effect of time (‘Post’ – ‘Pre’ intervention). Four tests were conducted: one to investigate the effects of motor training on acceleration, and one to investigate each of the effects of motor training, tACS and PAS on MEP amplitude. We used Cauchy priors, centred on zero, with default values in JASP; these priors assume a higher likelihood of small effect sizes relative to large effect sizes. In these analyses, the data are used to gather evidence for or against the null hypothesis (H_0_: no difference between measurements) or the alternative hypothesis (H_1_: existence of a difference between measurements; Wagenmakers et al. 2018), which is expressed using a Bayes factor. We report BF_10_, to quantify evidence favouring the alternative hypothesis over the null, for t-tests, and BF_incl_, the Bayes factor for inclusion of an effect, calculated across all models containing that effect within an ANOVA model. Effect sizes are reported as partial eta-squared (for ANOVA effects) or as Cohen’s d (for t-tests). We adopted the following labelling convention: BF_10_ >10 indicates strong support for H_1_ over H_0_; BF_10_ >3 indicates moderate support for H_1_ over H_0_; 1< BF <3 indicates anecdotal support for H_1_ over H_0_; 1/3< BF_10_ <1 indicates anecdotal support for H_0_ over H_1_; 1/10< BF_10_ <1/3 indicates moderate support for H_0_ over H_1_; BF_10_ <1/10 indicates strong evidence for H_0_ over H_1_; and BF_10_ = 1 indicates no evidence for H_0_ or H_1_.

## Results

### Motor training induces robust increases in performance and MEPs

In the first phase of each session, participants performed 30 minutes of ballistic abduction of the *abductor pollicis brevis* (APB), which we hypothesised would increase acceleration and MEP amplitudes in all conditions. For active and sham tACS conditions, acceleration was higher after motor training than at baseline (**Table 1**). A Bayesian two-way repeated-measures ANOVA on horizontal thumb acceleration before and after motor training revealed strong evidence for a main effect of time (BF_incl_ = ∞, η_p_^2^ = 0.74), with moderate evidence against a main effect of tACS type (BF_incl_ = 0.2, η_p_^2^ = 0.008) and moderate evidence against their interaction (BF_incl_ = 0.3, η_p_^2^ = 0.056).

**Table 1.** Acceleration (arbitrary units). Mean ± SD

|  | N | Pre MT | Post MT |
| --- | --- | --- | --- |
| Active tACS | 36 | 1.51 ± 0.7 | 2.32 ± 0.7 |
| Sham tACS | 36 | 1.36 ± 0.63 | 2.37 ± 0.56 |
| Control tACS | 38 | 1.44 ± 0.55 | 1.95 ± 0.53 |

Consistent with our hypothesis and with previous results using the same or similar training paradigms, motor training resulted in an increase in MEP amplitude (**Fig. 2A, 2B** left panel). A Bayesian two-way repeated-measures ANOVA of MEP amplitude revealed strong evidence for a main effect of motor training (BF_incl_ = 879.2, η_p_^2^ = 0.249), with moderate evidence against a main effect of tACS type (BF_incl_ = 0.14, η_p_^2^ = 8.55 x e^-4^) and moderate evidence against their interaction (BF_incl_ = 0.15, η_p_^2^ = 0.012), reflecting larger MEP amplitudes after motor training than at baseline. For the control tACS condition, both acceleration and MEPs were increased after motor training (**Fig. 3A, 3B** left panel). A Bayesian paired t-test on horizontal thumb acceleration revealed strong evidence that acceleration differed before and after training (BF_10_ = 6.27 x e^5^, Cohen’s d = 1.02). Similarly, there was moderate evidence that MEP amplitudes differed before and after training (BF_10_ = 6.73, Cohen’s d = 0.48). These results confirm that the initial plasticity paradigm (motor training by ballistic thumb abduction) was effective in improving performance and in inducing plasticity in the motor system.

**Figure 2.**
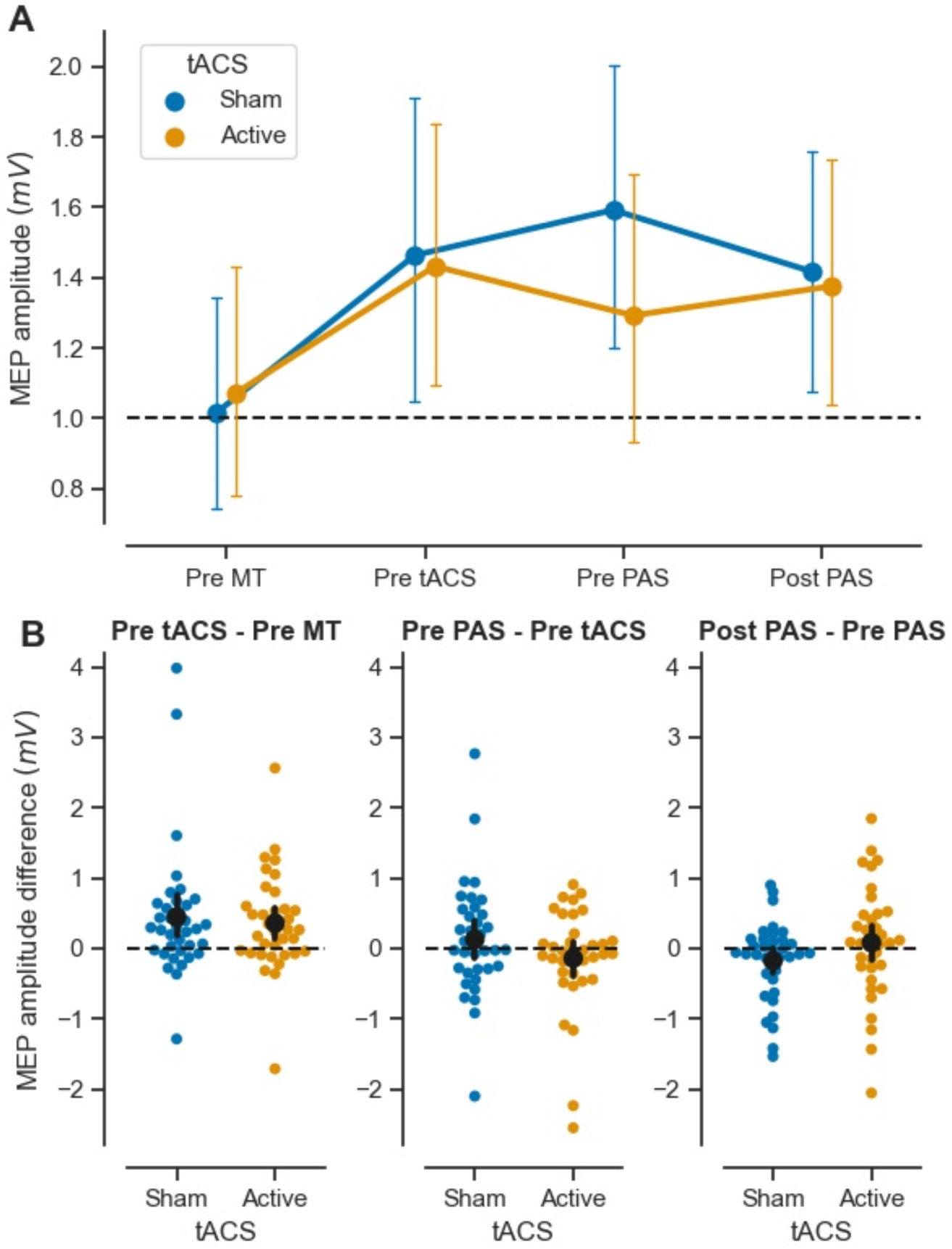
Effect of sham and active tACS on MEP amplitude in response to successive plasticity paradigms. **A)** Raw MEP amplitude (mV) at four experimental time-points: at baseline (Pre MT), after motor training/before tACS (Pre tACS), after tACS/before PAS (Pre PAS) and after PAS (Post PAS), in the sham (blue) and active (orange) tACS sessions. Symbols represent the mean; error bars denote 95% confidence intervals (CI, 1000 bootstrap iterations). Dotted line denotes baseline MEP amplitude. **B)** Mean MEP amplitude difference over each intervention (Post – Pre, black symbol), with 95% CI (black bar) and individual observations (coloured symbols). Effect of 30 minutes of ballistic motor training (MT) on MEP amplitude (left). Note that ‘active’ and ‘sham’ refer to active and sham tACS sessions and that no brain stimulation had been delivered yet. Effect of 18 min. of slow-oscillatory transcranial alternating current stimulation (tACS) (middle). Effect of paired associative stimulation (PAS_25_, excitatory protocol) on MEP amplitude (right). Dotted line denotes null difference between time points. N=36.

**Figure 3.**
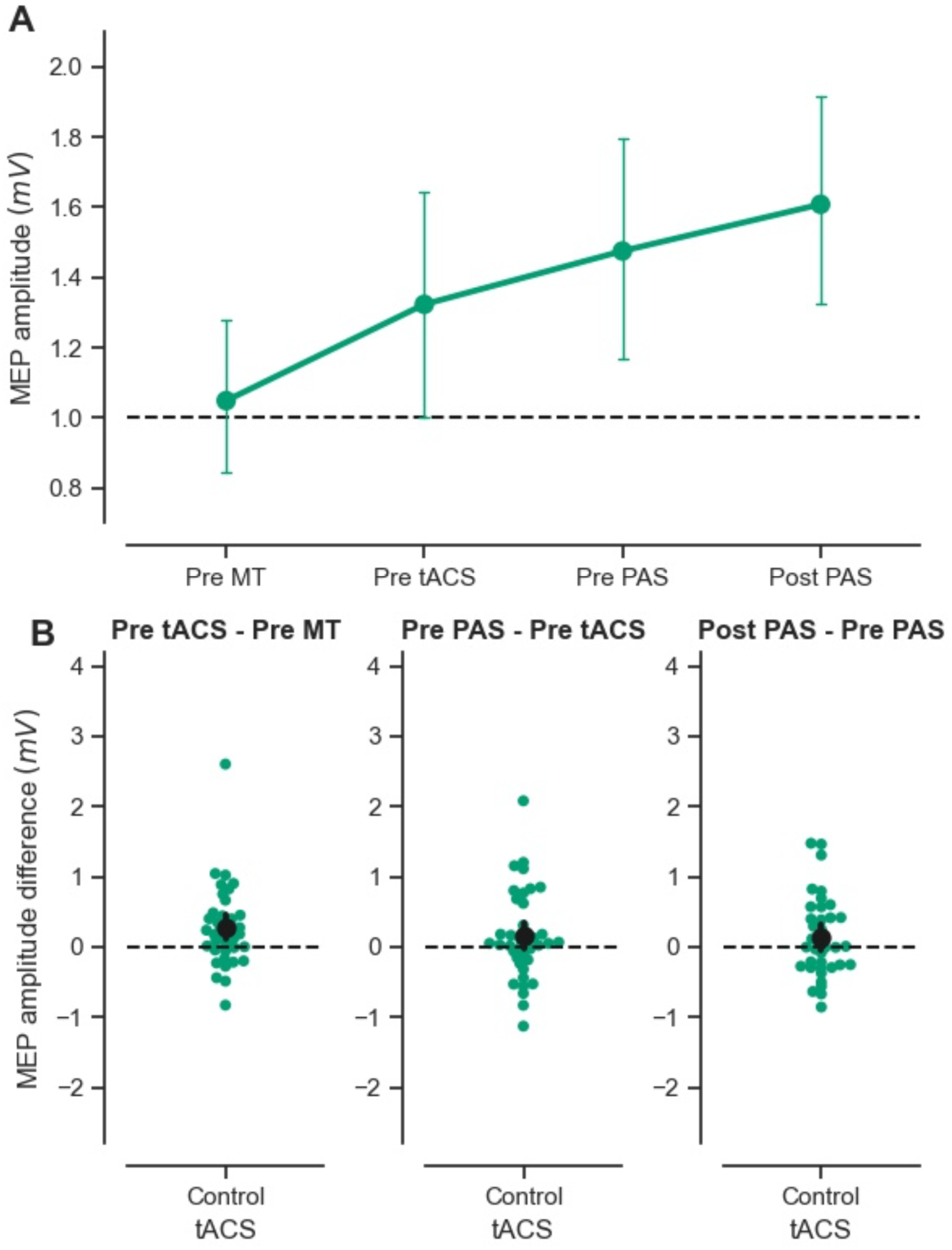
Effect of control tACS on MEP amplitude in response to successive plasticity paradigms. **A)** Raw MEP amplitude (mV) for the four experimental time-points: at baseline (Pre MT), after motor training/before tACS (Pre tACS), after tACS/before PAS (Pre PAS) and after PAS (Post PAS), in the control (green) tACS sessions Symbols represent the mean; error bars denote 95% confidence intervals (CI, 1000 bootstrap iterations). Dotted line denotes baseline MEP amplitude. **B)** Mean MEP amplitude difference over each intervention (Post – Pre, black symbol), with 95% CI (black bar) and individual observations (coloured symbols). Effect of 30 minutes of ballistic motor training (MT) on MEP amplitude (left). Effect of 18 min. of slow-oscillatory transcranial alternating current stimulation (tACS) (middle). Effect of paired associative stimulation (PAS_25_, excitatory protocol) on MEP amplitude (right). Dotted line denotes null difference between time points. N=38.

### tACS following motor training: acute effects on MEPs

Next, we examined the effect of tACS on MEPs immediately after tACS, and prior to PAS. We hypothesised that in the sham tACS and control tACS conditions, MEPs would remain elevated relative to baseline, or increase further as a consequence of motor learning. We did not have a specific hypothesis concerning MEPs following active tACS, as slow oscillations could be hypothesised to reduce MEPs, or to modulate synaptic scaling without affecting MEP amplitude directly. Overall, there was little evidence of further change in MEP amplitudes after tACS (**Fig. 2B, 3B**, middle panels), consistent with previous research demonstrating that MEP increases induced by motor training persist for at least 30 minutes following training (Ziemann et al., 2004). For the sham and active tACS conditions, a Bayesian two-way repeated-measures ANOVA revealed moderate evidence against a main effect of time (BF_incl_ = 0.14, η_p_^2^ = 7.46 x e^-5^), anecdotal evidence against a main effect of tACS type (BF_incl_ = 0.37, η_p_^2^ = 0.039) and strong evidence against the interaction of time and tACS type (BF_incl_ = 0.1, η_p_^2^ = 0.118). For the control tACS condition, a Bayesian paired t-test before and after tACS provided anecdotal evidence that MEP amplitudes did not differ over time (BF_10_ = 0.47, Cohen’s d = 0.24).

### tACS following motor training does not enable subsequent PAS plasticity

A prediction arising from the synaptic homeostasis hypothesis is that slow wave oscillations “reset” synaptic connections back to a functional range, making them more receptive to subsequent plasticity paradigms. Thus, we predicted that the excitatory effect of PAS should manifest more robustly when preceded by active tACS (which should “unload” the targeted synapses) compared with sham tACS (where synaptic plasticity should have become relatively more saturated). In contrast, the control tACS montage should not “reset” synaptic connections in the network of interest and thus should not prevent interference of motor training with PAS. We found evidence suggesting that active tACS does not restore PAS effects after motor training (**Fig. 2B** right panel). Furthermore, results were equivocal with respect to the question of whether PAS modulates MEP amplitudes after sham tACS and control tACS (**Fig. 2B, 3B** right panels). For the sham and active tACS conditions, a Bayesian two-way repeated-measures ANOVA revealed moderate evidence against a main effect of time (BF_incl_ = 0.15, η_p_^2^ = 0.007), anecdotal evidence against a main effect of tACS (BF_incl_ = 0.43, η_p_^2^ = 0.042) and moderate evidence against the interaction of time and tACS type (BF_incl_ = 0.11, η_p_^2^ = 0.084). Post-hoc Bayesian paired t-tests revealed anecdotal and moderate evidence levels against an effect of PAS following sham tACS (BF_10_ = 0.88, Cohen’s d = -0.32) and active tACS (BF_10_ = 0.22, Cohen’s d = 0.1), respectively. For the control tACS condition, a Bayesian paired t-test revealed anecdotal evidence against the effect of time on MEP amplitudes (BF_10_ = 0.46, Cohen’s d = 0.24).

Exploratory visualisation of the relationship between the effect of different interventions (expressed as the difference in MEP amplitudes Post – Pre intervention) did not reveal any trends in the data beyond those outlined above (**Fig. 4**).

**Figure 4.**
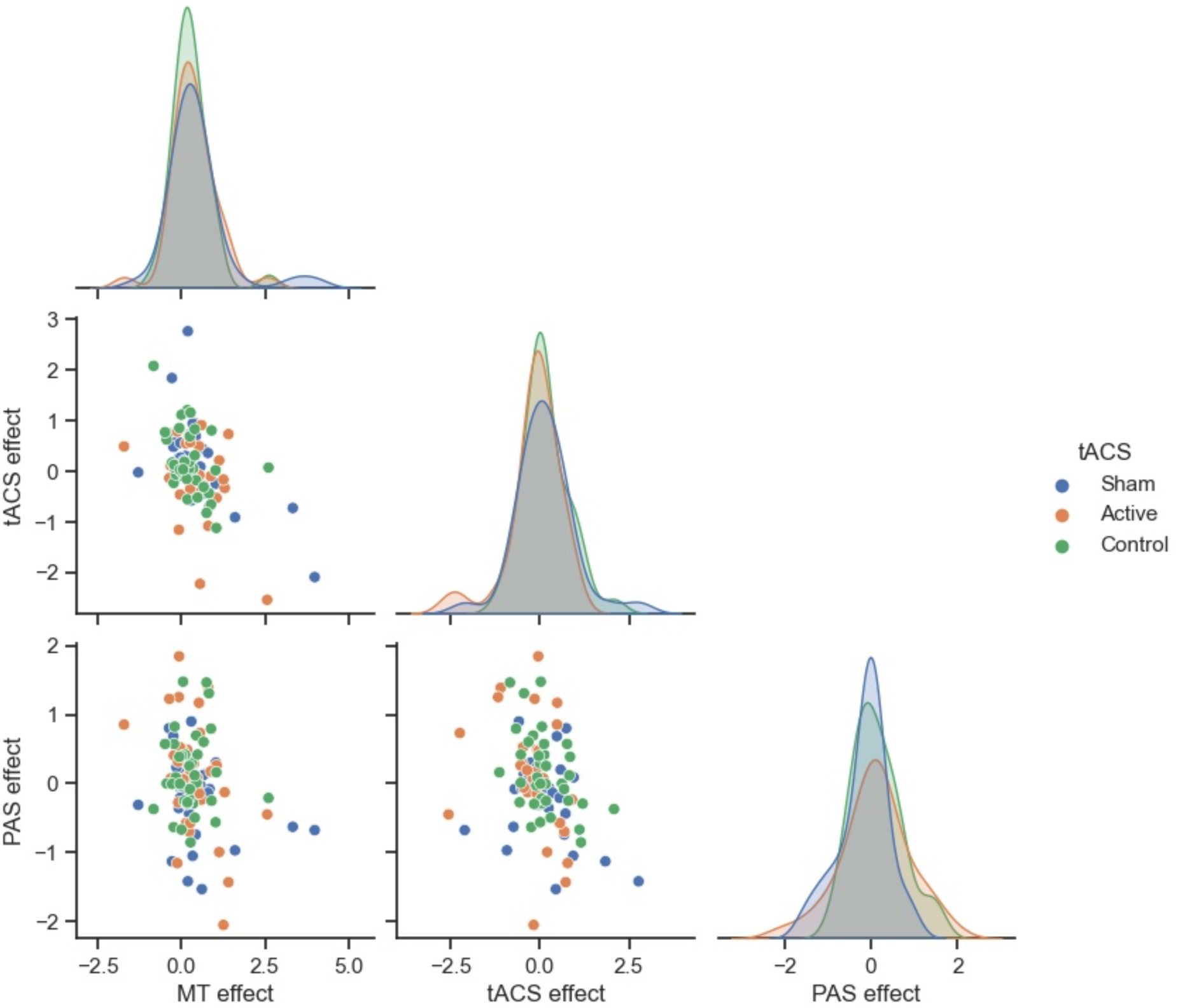
Relationship between the effects of successive plasticity interventions. Scatterplots comparing the effect of motor training (MT), slow-oscillatory transcranial alternating current stimulation (tACS), and paired associative stimulation (PAS), in the sham fronto-motor (blue), active fronto-motor (orange) and control posterior midline (green) tACS conditions. Results are expressed as the difference in MEP amplitude (mV) (Post – Pre) for the different interventions. Symbols represent individuals and shaded curves represent estimates of the probability density functions of the variables. Regions of overlap between the curves are shown in grey.

### Assessing PAS effects using an adjusted stimulation intensity

In a subset of participants who received active and sham tACS (N=29), intensity of stimulation was readjusted prior to PAS to elicit MEPs of similar amplitude to baseline. This new intensity was used to record MEPs before and after PAS to address potential concerns that PAS effects may be masked by ceiling effects. Results did not provide conclusive evidence that active tACS modulates PAS effects differently to sham tACS (**Table 2**). A Bayesian two-way repeated-measures ANOVA revealed moderate evidence in favour of a main effect of time (BF_incl_ = .062, η_p_^2^ = 0.251), anecdotal evidence against a main effect of tACS (BF_incl_ = 0.73, η_p_^2^ = 0.069) and anecdotal evidence against the interaction of time and tACS type (BF_incl_ = 0.49, η_p_^2^ = 0.024). Post-hoc Bayesian paired t-tests revealed anecdotal and moderate evidence for an effect of PAS following sham tACS (BF_10_ = 1.16, Cohen’s d = 0.38) and active tACS (BF_10_ = 3.53, Cohen’s d = 0.49), respectively. However, there was anecdotal evidence against a difference in MEP amplitudes between active and sham sessions after PAS (BF_10_ = 0.59, Cohen’s d = 0.29).

**Table 2.**
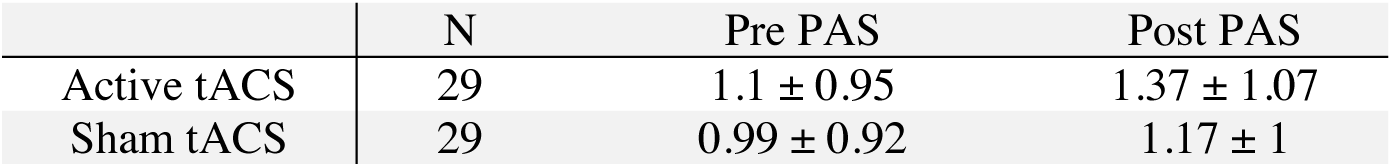
MEP amplitude (mV), recorded at adjusted intensity. Mean ± SD

### No evidence for specific entrainment of slow-oscillations immediately after active tACS

To investigate whether active tACS entrained brain oscillations at its frequency of delivery (0.75 Hz) or caused other modulations of oscillatory activity, we investigated changes in scalp EEG power immediately after the three blocks of 6 minutes of tACS, as compared to just before (**Fig.5A**). A (non-Bayesian) cluster-based permutation test revealed no significant difference at baseline between sham and active tACS sessions. The same statistical approach uncovered a significant increase in power after both active (2 clusters: p=0.001, cluster statistic=320; p=0.006, cluster statistic=206) and sham (2 clusters: p=0.001, cluster statistic=413; p=0.009, cluster statistic=151) tACS separately (**Fig.5B**). Following both sham and active stimulation, there was a widespread power increase in slow-oscillation (0.5-1Hz) and delta (1-4Hz) frequency bands, as well as a more spatially limited increase in high-alpha (12-15Hz) and beta (15-25Hz) frequencies. Direct comparison of both raw power and decibel-change values from baseline revealed no significant difference between active and sham sessions post-tACS. Focusing on the tACS frequency band, a two-way repeated measures ANOVA of pooled power values in the electrodes surrounding the motor tACS pad revealed a significant effect of time (F(3,72)=7.9, p=0.0001), but not of tACS type (active or sham, F(1,24)=0.008, p=0.93), and there was no interaction between these factors (F(3,72)=0.2, p=0.89) (**Fig.5C**). Thus, there was no evidence for a frequency-specific entrainment effect in the EEG data that outlasted the period of direct stimulation.

**Figure 5.**
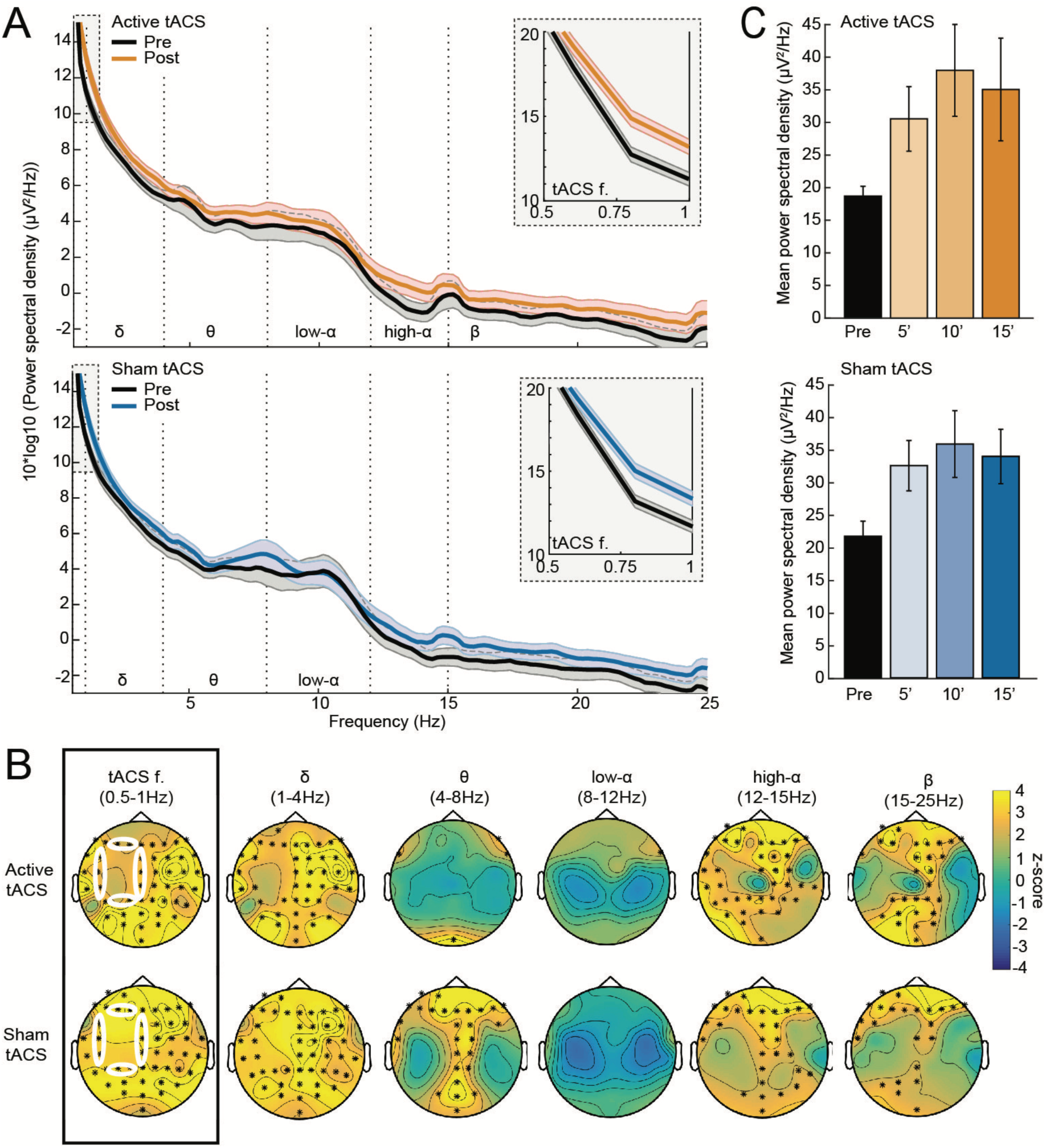
Off-line modulation of EEG power by tACS. **A)** Average EEG power spectrum in the electrodes surrounding the motor tACS pad (10 pooled electrodes, see white circles and details in panel B), just before tACS (black curves) and immediately after each block of tACS (average of three 1-minute segments, coloured curves: active tACS in orange, sham tACS in blue). The tACS frequency band (0.5-1Hz) is highlighted in grey and enlarged in the grey inset for readability. Error bars denote SEM, N=25 participants. **B)** Cluster-based permutation analysis contrasting power values before and after tACS (top row: active, bottom row: sham) in each of the 6 frequency bands of interest (tACS frequency, delta, theta, low-alpha, high-alpha and beta). Scalp maps display z-scores of the initial sample statistic and * denotes p< 0.01 for electrodes belonging to a significant cluster. The 10 electrodes surrounding the motor tACS pad (FCz, Cz, CPz, CP5, C5, FC5, F3, P3, F1, P1) are highlighted by white ellipses in the scalp maps of tACS frequency (black rectangle). **C)** Mean power in the tACS frequency band before, after 5 minutes, 10 minutes and 15 minutes of tACS, for active tACS (top, orange) and sham tACS (bottom, blue). Average across participants in the pooled electrodes surrounding the motor tACS pad. An ANOVA revealed a significant effect of time, but there was no effect of stimulation type or interaction between these factors. Errors bars denote SEM, N=25 participants.

## Discussion

We set out to test whether a short period of tACS could attenuate the interference between two successive motor plasticity paradigms in awake humans. We found that motor training by repetitive, ballistic thumb movements induced an increase in MEPs. Importantly, we found moderate evidence against an effect of active tACS in restoring PAS plasticity, together with no evidence of lasting entrainment of slow-oscillations in the EEG. This suggests that, under the conditions tested here, slow-oscillatory tACS does not modulate synaptic homeostasis in the motor system of awake humans.

As some properties of PAS are reminiscent of long-term potentiation (LTP) and long-term depression (LTD) properties in animal preparations (e.g., NMDA-receptor dependence, time-course, spatial specificity) (Stefan et al., 2000, 2002; Wolters et al., 2003), PAS effects in humans have been labelled as ‘LTP/LTD-like’ plasticity (Stefan et al., 2000; Wolters et al., 2003; Ziemann et al., 2004) and interpreted as being analogous to neuronal LTP/LTD in animal experiments. The fact that PAS effects are impeded or even reversed by prior motor practice suggests that the initial motor practice triggered LTP-like plasticity, leading to modification of the LTP/LTD induction threshold, and tipping the balance in favour of LTD-like plasticity induction (Ziemann et al., 2004). Here, we showed attenuation but not reversal of ‘excitatory’ PAS effects by motor practice in the sham and control conditions, which is compatible with the notions of saturation of LTP-like processes and modification of the LTP/LTD threshold. Importantly, following active tACS, PAS did not lead to an enhancement of MEPs that was substantially different from sham. This could have occurred for a number of reasons, which are discussed below.

While there is no consensus regarding the mechanisms of action of tACS, recent literature has focused on detecting local, long-lasting increases in oscillatory power at the stimulation frequency, which could be explained in terms of entrainment of cortical oscillatory activity (Ozen et al., 2010; Ali et al., 2013; Helfrich et al., 2014; Alagapan et al., 2016; although see Vossen et al., 2015 for a network plasticity account). More specifically, with regard to slow oscillations, a number of studies using scalp EEG recordings in humans have reported locally increased power in the low-frequency band immediately following slow-oscillatory tDCS to frontal cortex (Marshall et al., 2006; Kirov et al., 2009; Antonenko et al., 2013; Westerberg et al., 2015; Paßmann et al., 2016; Ladenbauer et al., 2016, 2017). In contrast, the current study found no evidence for a specific power increase in the low-frequency band immediately following active tACS.

This apparent discrepancy with previous studies is likely attributable to several factors. First, all but one previous study delivered slow-oscillatory tDCS during sleep (either a nap or a full-night’s sleep; Marshall et al., 2006; Antonenko et al., 2013; Westerberg et al., 2015; Paßmann et al., 2016; Ladenbauer et al., 2016, 2017), when cortical slow-oscillations are expected to occur endogenously (Steriade et al., 1993). It is possible that to achieve modulation of slow oscillations via tACS, the network needs to be in a state that enables generation of slow oscillations, or that such oscillations need to be present at the time of stimulation. Consistent with this hypothesis, both modelling and animal stimulation experiments have shown that weak oscillatory currents are most effective at modulating endogenous oscillations in cortical networks (Schmidt et al., 2014). Second, most previous studies employed a stimulation waveform containing a direct current (DC) component (slow-oscillatory tDCS), as opposed to the pure tACS used here. Even though such an explanation would render frequency-specific power modulations harder to account for, it is conceivable that the EEG after-effects described in previous literature resulted from the DC component rather than the oscillating component of slow-oscillatory tDCS. Third, in previous studies, stimulation was applied to the frontal cortex, which has been shown to be a prominent source of slow oscillations (Massimini et al., 2004; Sheroziya et al., 2014). It is conceivable that tACS applied through a classic ‘motor’ parieto-frontal montage does not modulate low-frequency oscillations because it targets a fundamentally different functional network that is less reliant on slow oscillations to achieve consolidation of memories. However, studies have highlighted the existence of ‘local’ slow-oscillations, even in motor cortex (Huber et al., 2013; Fattinger et al., 2017). Fourth, as our evaluation of tACS effects on EEG took place immediately after an episode of motor learning, it is possible that learning-related EEG modulations in both the sham and active tACS conditions masked the effects of tACS alone. Finally, it is important to note that a number of studies using variations of slow-oscillatory tDCS/tACS failed to find lasting modulation of low-frequency oscillations (Eggert et al., 2013; Sahlem et al., 2015; D’Atri et al., 2016; Bueno-Lopez et al., 2019). Most notably, Lafon et al. (2017) reported a lack of entrainment to slow-wave tACS, as measured by implanted electrodes in human epileptic patients.

Importantly, the EEG results reported here do not speak to the likelihood of other mechanisms of action of tACS (for a review, see Liu et al., 2018), such as modulation of neuronal activity and excitability *during* tACS. We recorded EEG during tACS, but did not analyse the resulting data, as the physiological electric signal is strongly distorted by the tACS-injected current, and satisfactory correction of these distortions is still very much under debate (see e.g., Noury et al., 2017; Neuling et al., 2017). Other hypothesised neuronal mechanisms of action are currently the subject of intense discussion, with some authors suggesting that the stimulation intensities currently used might not be sufficient to cause measurable modulation of neuronal activity (Lafon et al. 2017; Vöröslakos et al. 2018, but see Krause et al. 2019). Interestingly, a recent study highlighted the possibility of *indirect* neural entrainment at a cortical level via the action of tACS on peripheral nerves (Asamoah et al. 2019).

The current study has a number of limitations. Additional experiments assessing the effect of tACS alone and tACS following motor training (with no subsequent PAS) on MEP amplitude could help ascertain whether tACS has MEP-modulating effects on its own, and if so, whether it interacts with motor training in a homeostatic way. However, previous research has failed to show any effects of slow oscillatory tACS on motor cortical excitability (Antal et al., 2008). We did not re-test ballistic thumb movement acceleration after tACS or after PAS. Consequently, we are not able to ascertain whether the levels of performance were maintained throughout the session. Our motivation for not re-assessing motor performance was two-fold. First, we wished to avoid any interference that might arise from having participants perform the motor task again, as it has been suggested that motor activity can reverse or abolish theta-burst stimulation-induced changes (Iezzi et al., 2008). Second, additional thumb acceleration measurements would have added time to an already lengthy experiment, which might have adversely impacted participant arousal level or motivation.

In conclusion, we interleaved tACS between two interventions known to induce plasticity in human motor cortex, and which have been shown previously to interfere with each other. We found evidence against a modulation of cortico-spinal excitability by PAS after active tACS, which suggests that under the conditions used here, tACS does not modulate plasticity interactions in the motor cortex.

## Acknowledgments

This work was supported by a grant from the National Health and Medical Research Council of Australia (APP1078464) awarded to MVS and JBM. MVS and JBM were also supported by an Office of Naval Research Global grant (N62909-17-1-2139). JBM was supported by an Australian Research Council (ARC) Australian Laureate Fellowship (FL110100103) and the ARC Centre of Excellence for Integrative Brain Function (ARC Centre Grant CE140100007). None of the funding sources took part in study design; in the collection, analysis and interpretation of data; in the writing of the report; or in the decision to submit the article for publication. The authors thank Jessica Lister for assistance with data collection and David Lloyd with graphical illustration of the experimental setup.

## References

Abraham WC, Bear MF (1996) Metaplasticity : plasticity of synaptic. Trends Neurosci 19:126–130.

Aeschbach D, Cutler AJ, Ronda JM (2008) A Role for Non-Rapid-Eye-Movement Sleep Homeostasis in Perceptual Learning. J Neurosci 28:2766–2772.

Alagapan S, Schmidt SL, Levebvre J, Hadar E, Schin HW, Frohlich F (2016) Modulation of Cortical Oscillations by Low-Frequency Direct Cortical Stimulation Is State-Dependent. PLoS Biol 14:e1002424.

Ali MM, Sellers KK, Frohlich F (2013) Transcranial Alternating Current Stimulation Modulates Large-Scale Cortical Network Activity by Network Resonance. J Neurosci 33:11262–11275.

Antal A, Boros K, Poreisz C, Chaieb L, Terney D, Paulus W (2008) Comparatively weak after-effects of transcranial alternating current stimulation (tACS) on cortical excitability in humans. Brain Stimul 1:97–105.

Antonenko D, Diekelmann S, Olsen C, Born J, Mölle M (2013) Napping to renew learning capacity: Enhanced encoding after stimulation of sleep slow oscillations. Eur J Neurosci 37:1142–1151.

Asamoah B, Khatoun A, Mc Laughlin M. (2019) tACS motor system effects can be caused by transcutaneous stimulation of peripheral nerves. Nat Commun. 10(1):266.

Biel AL, Friedrich EVC (2018) Why You Should Report Bayes Factors in Your Transcranial Brain Stimulation Studies. Front Psychol. 2;9:1125.

Bueno-Lopez A, Eggert T, Dorn H, Danker-Hopfe H (2019) Slow oscillatory transcranial direct current stimulation (so-tDCS) during slow wave sleep has no effects on declarative memory in healthy young subjects. Brain Stimul 12:948–958.

D’Atri A, De Simoni E, Gorgoni M, Ferrara M, Ferlazzo F, Rossini PM, De Gennaro L (2016) Electrical stimulation of the frontal cortex enhances slow-frequency EEG activity and sleepiness. Neuroscience 324:119–130.

Delvendahl I, Jung NH, Kuhnke NG, Ziemann U, Mall V (2012) Plasticity of motor threshold and motor-evoked potential amplitude--a model of intrinsic and synaptic plasticity in human motor cortex? Brain Stimul ;5(4):586–93.

Dienes Z, Mclatchie N (2018) Four reasons to prefer Bayesian analyses over significance testing. Psychon Bull Rev ;25(1):207–218.

Diekelmann S, Born J (2010) The memory function of sleep. Nat Rev Neurosci 11:114–26.

Eggert T, Dorn H, Sauter C, Nitsche MA, Bajbouj M, Danker-Hopfe H (2013) No effects of slow oscillatory transcranial direct current stimulation (tDCS) on sleep-dependent memory consolidation in healthy elderly subjects. Brain Stimul 6:938–945.

Fattinger S, de Beukelaar TT, Ruddy KL, Volk C, Heyse NC, Herbst JA, Hahnloser RHR, Wenderoth N, Huber R (2017) Deep sleep maintains learning efficiency of the human brain. Nat Commun 8:15405.

Helfrich RF, Schneider TR, Rach S, Trautmann-Lengsfeld SA, Engel AK, Herrmann CS (2014) Entrainment of brain oscillations by transcranial alternating current stimulation. Curr Biol 24:333–339.

Hodgson RA, Ji Z, Standish S, Boyd-Hodgson TE, Henderson AK, Racine RJ (2005) Training-induced and electrically induced potentiation in the neocortex. Neurobiol. Learn Mem 83:22–32.

Huber R, Maki H, Rosanova M, Casarotto S, Canali P, Casali AG, Tononi G, Massimini M (2013) Human cortical excitability increases with time awake. Cereb. Cortex 23:332–338.

Iezzi E, Conte A, Suppa A, Agostino R, Dinapoli L, Scontrini A, Berardelli A (2008) Phasic Voluntary Movements Reverse the Aftereffects of Subsequent Theta-Burst Stimulation in Humans. J Neurophysiol 100:2070–2076.

JASP Team (2020). JASP (Version 0.14.1) [Computer software].

Kamke MR, Ryan AE, Sale MV, Campbell ME, Riek S, Carroll TJ, Mattingley JB (2014) Visual Spatial Attention Has Opposite Effects on Bidirectional Plasticity in the Human Motor Cortex. J Neurosci 34:1475–1480.

Keel JC, Smith MJ, Wassermann EM (2000) A safety screening questionnaire for transcranial magnetic stimulation. Clin Neurophysiol 112:720.

Kirov R, Weiss C, Siebner HR, Born J, Marshall L (2009) Slow oscillation electrical brain stimulation during waking promotes EEG theta activity and memory encoding. Proc Natl Acad Sci 106:15460–15465.

Krause MR, Vieira PG, Csorba BA, Pilly PK, Pack CC. (2019) Transcranial alternating current stimulation entrains single-neuron activity in the primate brain. Proc Natl Acad Sci U S A. 116(12):5747–5755.

Kuhn M, et al. (2016) Sleep recalibrates homeostatic and associative synaptic plasticity in the human cortex. Nat Commun 7:12455.

Ladenbauer J, Kulzow N, Passmann S, Antonenko D, Grittner U, Tamm S, Floel A (2016) Brain stimulation during an afternoon nap boosts slow oscillatory activity and memory consolidation in older adults. Neuroimage 142:311–323.

Ladenbauer J, Ladenbauer J, Kulzow N, de Boor R, Avramova E, Grittner U, Floel A (2017) Promoting Sleep Oscillations and Their Functional Coupling by Transcranial Stimulation Enhances Memory Consolidation in Mild Cognitive Impairment. J Neurosci 37:7111–7124.

Lahr J, Paßmann S, List J, Vach W, Flöel A, Klöppel S (2016) Effects of Different Analysis Strategies on Paired Associative Stimulation. A Pooled Data Analysis from Three Research Labs. PLoS One 2016 May 4;11(5):e0154880.

Lafon B, Henin S, Huang Y, Friedman D, Melloni L, Thesen T, Doyle W, Buzsaki G, Devinsky O, Parra LC, Liu A (2017) Low frequency transcranial electrical stimulation does not entrain sleep rhythms measured by human intracranial recordings. Nat Commun 8:1199.

Liu A, Voroslakos M, Kronberg G, Henin S, Krause MR, Huang Y, Opitz A, Mehta A, Pack CC, Krekelberg B, Berenyi A, Parra LC, Melloni L, Devinsky O, Buzsaki G (2018) Immediate neurophysiological effects of transcranial electrical stimulation. Nat Commun 9:5092.

Mander BA, Santhanam S, Saletin JM, Walker MP (2011) Wake deterioration and sleep restoration of human learning. Curr Biol 21:R183–R184.

Maquet P (2001) The role of sleep in learning and memory. Science. 294:1048–1052.

Marshall L, Helgadóttir H, Mölle M, Born J (2006) Boosting slow oscillations during sleep potentiates memory. Nature 444:610–613.

Mascetti L, et al. (2013) The Impact of Visual Perceptual Learning on Sleep and Local Slow-Wave Initiation. J Neurosci 33:3323–3331.

Massimini M, Huber R, Ferrarelli F, Hill S, Tononi G (2004) The Sleep Slow Oscillation as a Traveling Wave. J Neurosci 24:6862–6870.

Mölle M, Marshall L, Gais S, Born J (2004) Learning increases human electroencephalographic coherence during subsequent slow sleep oscillations. Proc Natl Acad Sci U S A 101:13963–13968.

Moser EI, Krobert KA, Moser MB, Morris RGM (1998) Impaired spatial learning after saturation of long-term potentiation. Science 281:2038–2042.

Muellbacher W, Ziemann U, Wissel J, Dang N, Kofler M, Facchini S, Boroojerdi B, Poewe W, Hallett M (2002) Early consolidation in human primary motor cortex. Nature 415:640–644.

Muellbacher W, Ziemann U, Boroojerdi B, Cohen L, Hallett M (2001) Role of the human motor cortex in rapid motor learning. Exp Brain Res 136:431–438.

Neuling T, Ruhnau P, Weisz N, Herrmann CS, Demarchi G (2017) Faith and oscillations recovered: On analyzing EEG/MEG signals during tACS. Neuroimage 147:960–963.

Noury N, Siegel M (2017) Phase properties of transcranial electrical stimulation artifacts in electrophysiological recordings. Neuroimage 158:406–416.

Oldfield R C (1971) The assessment and analysis of handedness: The Edinburgh inventory. Neuropsychologia, 9, 97–113.

Ozen S, Sirota A, Belluscio MA, Anastassiou CA, Stark E, Koch C, Buzsaki G (2010) Transcranial Electric Stimulation Entrains Cortical Neuronal Populations in Rats. J Neurosci 30:11476–11485.

Paßmann S, Kulzow N, Ladenbauer J, Antonenko D, Grittner U, Tamm S, Floel A (2016) Boosting Slow Oscillatory Activity Using tDCS during Early Nocturnal Slow Wave Sleep Does Not Improve Memory Consolidation in Healthy Older Adults. Brain Stimul 9:730–739.

Ridding MC, Ziemann U (2010) Determinants of the induction of cortical plasticity by non-invasive brain stimulation in healthy subjects. J Physiol 588:2291–2304.

Rioult-Pedotti MS, Friedman D, Donoghue JP (2000) Learning-induced LTP in neocortex. Science 290:533–536.

Rossini PM, et al. (1994) Non-invasive electrical and magnetic stimulation of the brain, spinal cord and roots: basic principles and procedures for routine clinical application. Report of an IFCN committee. Electroencephalogr Clin Neurophysiol 91:79–92.

Rossini PM et al. (2015) Non-invasive electrical and magnetic stimulation of the brain, spinal cord, roots and peripheral nerves: Basic principles and procedures for routine clinical and research application. An updated report from an I.F.C.N. Committee. Clin Neurophysiol. ;126(6):1071–1107.

Sahlem GL, Badran BW, Halford JJ, Williams NR, Korte JE, Leslie K, Strachan M, Breedlove JL, Runion J, Bachman DL, Uhde TW, Borckardt JJ, George MS (2015) Oscillating square wave transcranial direct current stimulation (tDCS) delivered during slow wave sleep does not improve declarative memory more than sham: A randomized sham controlled crossover study. Brain Stimul 8:528–534.

Sale MV, Ridding MC, Nordstrom MA (2007) Factors influencing the magnitude and reproducibility of corticomotor excitability changes induced by paired associative stimulation. Exp Brain Res 181:615–626.

Sale MV, Ridding MC, Nordstrom MA (2008) Cortisol Inhibits Neuroplasticity Induction in Human Motor Cortex. J Neurosci 28:8285–8293.

Schmidt SL, Iyengar AK, Foulser AA, Boyle MR, Fröhlich F (2014) Endogenous cortical oscillations constrain neuromodulation by weak electric fields. Brain Stimul 7:878–889.

Shadmehr R, Brashers-Krug T (1997) Functional Stages in the Formation of Human Long-Term Motor Memory. J Neurosci 17:409–419.

Sheroziya M, Timofeev I (2014) Global Intracellular Slow-Wave Dynamics of the Thalamocortical System. J Neurosci 34:8875–8893.

Stefan K, Kunesch E, Cohen LG, Benecke R, Classen J (2000) Induction of plasticity in the human motor cortex by paired associative stimulation. Brain 123:572–84.

Stefan K, Kunesch E, Benecke R, Cohen LG, Classen J (2002) Mechanisms of enhancement of human motor cortex excitability induced by interventional paired associative stimulation. J Physiol 543:699–708.

Steriade M, McCormick DA, Sejnowski TJ (1993) Thalamocortical oscillations in the sleeping and aroused brain. Science 262:679–685.

Thielscher A, Antunes A, Saturnino GB (2015) Field modeling for transcranial magnetic stimulation: a useful tool to understand the physiological effects of TMS? IEEE EMBS 2015, Milano, Italy

Tononi G, Cirelli C (2014) Sleep and the Price of Plasticity: From Synaptic and Cellular Homeostasis to Memory Consolidation and Integration. Neuron 81:12–34.

Tucker MA, Hirota Y, Wamsley EJ, Lau H, Chaklader A, Fishbein W (2006) A daytime nap containing solely non-REM sleep enhances declarative but not procedural memory. Neurobiol Learn Mem 86:241–247.

Vöröslakos M, Takeuchi Y, Brinyiczki K, Zombori T, Oliva A, Fernández-Ruiz A, Kozák G, Kincses ZT, Iványi B, Buzsáki G, Berényi A. (2018) Direct effects of transcranial electric stimulation on brain circuits in rats and humans. Nat Commun. 9(1):483.

Vossen A, Gross J, Thut G (2015) Alpha power increase after transcranial alternating current stimulation at alpha frequency (a-tACS) reflects plastic changes rather than entrainment. Brain Stimul 8:499–508.

Wagenmakers, E.-J. et al. (2018). Bayesian inference for psychology, part I: theoretical advantages and practical ramifications. Psychon. Bull. Rev. 25, 35–37.

Westerberg CE, Florczak SM, Weintraub S, Mesulam MM, Marshall L, Zee PC, Paller KA (2015) Memory improvement via slow-oscillatory stimulation during sleep in older adults. Neurobiol Aging 36:2577–2586.

Wolters A, Sandbrink F, Schlottmann A, Kunesch E, Stefan K, Cohen LG, Benecke R, Classen J A (2003) Temporally Asymmetric Hebbian Rule Governing Plasticity in the Human Motor Cortex. J Neurophysiol 89:2339–2345.

Ziemann U, Iliac TV, Pauli C, Meintzschel F, Ruge D (2004) Learning Modifies Subsequent Induction of Long-Term Potentiation-Like and Long-Term Depression-Like Plasticity in Human Motor Cortex. J Neurosci 24:1666–1672.

Ziemann U, Siebner HR (2008) Modifying motor learning through gating and homeostatic metaplasticity. Brain Stimul 1:60–66.

